# CLEAR-ST: Physics-informed probabilistic decontamination of spatial transcriptomics by modeling mRNA lateral diffusion

**DOI:** 10.64898/2026.08.13.744615

**Authors:** Kun Ma, Yuanhua Huang, Joshua W.K. Ho

## Abstract

Spatial transcriptomics is a rapidly evolving technology that allows for the measurement of gene expression in a spatially resolved manner. However, one technical problem that occurs for many sequencing-based spatial transcriptomics platforms is the presence of mRNA lateral diffusion, where mRNA from one spot can bind to probes in another spot, leading to contamination and inaccurate gene expression measurements. In Visium-like assays, this artifact is often visible as structured out-of-tissue signal and boundary-associated expression halos, yet its magnitude, spatial decay, and directional bias vary substantially across samples. Here, we present CLEAR-ST, a physics-informed probabilistic framework for correcting diffusion-like contamination in spatial transcriptomics data. CLEAR-ST infers a latent clean expression field using a denoising autoencoder and links it to the observed counts through a graph-Laplacian forward contamination model with learnable diffusion parameters, finally evaluated with a selectable count likelihood. We first conducted a comprehensive comparison between 10X official and independently generated Visium samples, demonstrating that out-of-tissue count profiles are highly related to nearby in-tissue expression, more concentrated near tissue boundaries, and diffusion directions across genes are likely coherent. Across real samples with varying contamination burden, CLEAR-ST improved spatial domain recovery, increased gene-level spatial autocorrelation, and enhanced the biological specificity of downstream analyses such as marker gene discovery, pathway identification and cell type deconvolution. Compared to benchmark methods, CLEAR-ST showed consistent gains in clustering quality and concordance with manual annotations. Together, CLEAR-ST provides an interpretable and practical approach for diffusion-aware correction of capture-based spatial transcriptomics data.

## 1 Introduction

Spatial transcriptomics (ST) technologies measure gene expression while maintaining structural integrity of tissue, enabling the study of cellular neighborhoods, tissue architecture and organization, and disease associated microenvironments [1]. This technology has revolutionized our understanding of tissue organization and cellular interactions, providing insights into various biological processes and diseases [1]. Broadly speaking, there are two main types of spatial transcriptomics technologies: imaging-based and sequencing-based. Imaging-based technologies, such as MERFISH and seqFISH, use fluorescent probes to visualize mRNA molecules in situ, while sequencing-based technologies, such as 10x Genomics Visium and Slide-seq, capture mRNA from tissue sections and sequence them to obtain gene expression profiles. The latter has gained popularity due to its high throughput and ability to capture a large number of genes. However, the tissue permeabilization step must be sufficiently strong to release mRNA for capture by the slide, but excessive or spatially heterogeneous permeabilization can allow transcripts to spread beyond their original locations. This capture process introduces a fundamental vulnerability: transcripts may move away from their cells or tissue regions of origin before capture, thereby contaminating neighboring locations and distorting the measured spatial expression profile.

This phenomenon - lateral diffusion of mRNA, is a common technical issue in many sequencing-based spatial transcriptomics platforms [2] (Fig. 2a). Past studies have identified the issue experimentally. Through a human-mouse hybrid mRNA diffusion experiment, researchers discovered a significant amount of human transcripts in the mouse tissue spots and vice versa [3]. Another study compared across sequencing platforms to examine the extent of lateral diffusion [4]. Their results showed that tissue type and permeabilization time are the key factors in determining diffusion patterns: some tissues achieve optimal mRNA release with shorter duration (e.g., 6 minutes), while others require longer duration (e.g., 15 minutes). Even with the same sequencing platform, the extent of mRNA lateral diffusion could vary drastically, and hence it’s hard to conclude which platform is most resistant to the issue.

To mitigate the issue of mRNA lateral diffusion, there exist three categories of methods in the literature to the best of our knowledge. The first type of methods achieves denoising by solving it as a reference mapping or integration problem, and most often requires scRNA-seq data as reference [5, 6]. Another category focuses on removing noise signals, arguing that the complexity of diffusion patterns might lead to false positive with imputation [7]. Some representative ones include SpotGF and Split [7, 8]. The last type of methods models the diffusion process with some form of physical law or statistical framework to estimate the corrected expression profile. SpotClean models the diffusion process with a probabilistic framework and estimates the corrected expression profile, by modelling spot swapping and redistribute counts [3]. Another method resolVI uses a variational autoencoder to learn true expression by parameterizing the diffused amount from neighboring cells [9]. Two more recent approaches model diffusion reversion in a more physics-informed way - score matching, reminiscent of diffusion models in generative modeling [10, 11]. They model transcript locations instead of counts, and reversion is achieved by iteratively updating transcript locations with the learned score function. However, they are computationally intensive: each gene is fitted for its own score function and the reversion process requires multiple user-defined iterations. Hence, a new method that models the diffusion process with a physics-informed framework and is computationally efficient is still needed. We propose CLEAR-ST, a probabilistic decontamination framework that combines a denoising autoencoder with a spatial diffusion model to correct contamination in spatial transcriptomics data. The package is now available at https://github.com/holab-hku/CLEAR-ST.

## 2 Results

### 2.1 Comparative analysis of public and independently generated Visium datasets identifies diffusion-associated spatial patterns and motivates CLEAR-ST design

To characterize diffusion-associated signal and motivate the design of CLEAR-ST, we compared two collections of Visium data: 77 public Visium samples released by 10x Genomics and 14 independently generated Visium samples from a published study [12]. In Fig. 1a, we show one representative slide from each dataset and their unsupervised clustering results of full-slide spots (left), total count distribution in out-of-tissue spots (middle), and tissue boundary analysis within 5 grid distances (right). We observed that out-of-tissue spots frequently co-clustered with subsets of in-tissue spots, suggesting that a component of the background signal is spatially structured rather than uniformly distributed. This observation motivated our use of expression-derived diffusion-rate labels, which allow out-of-tissue spots to be linked to transcriptionally similar in-tissue regions during model initialization.

**Fig. 1.**
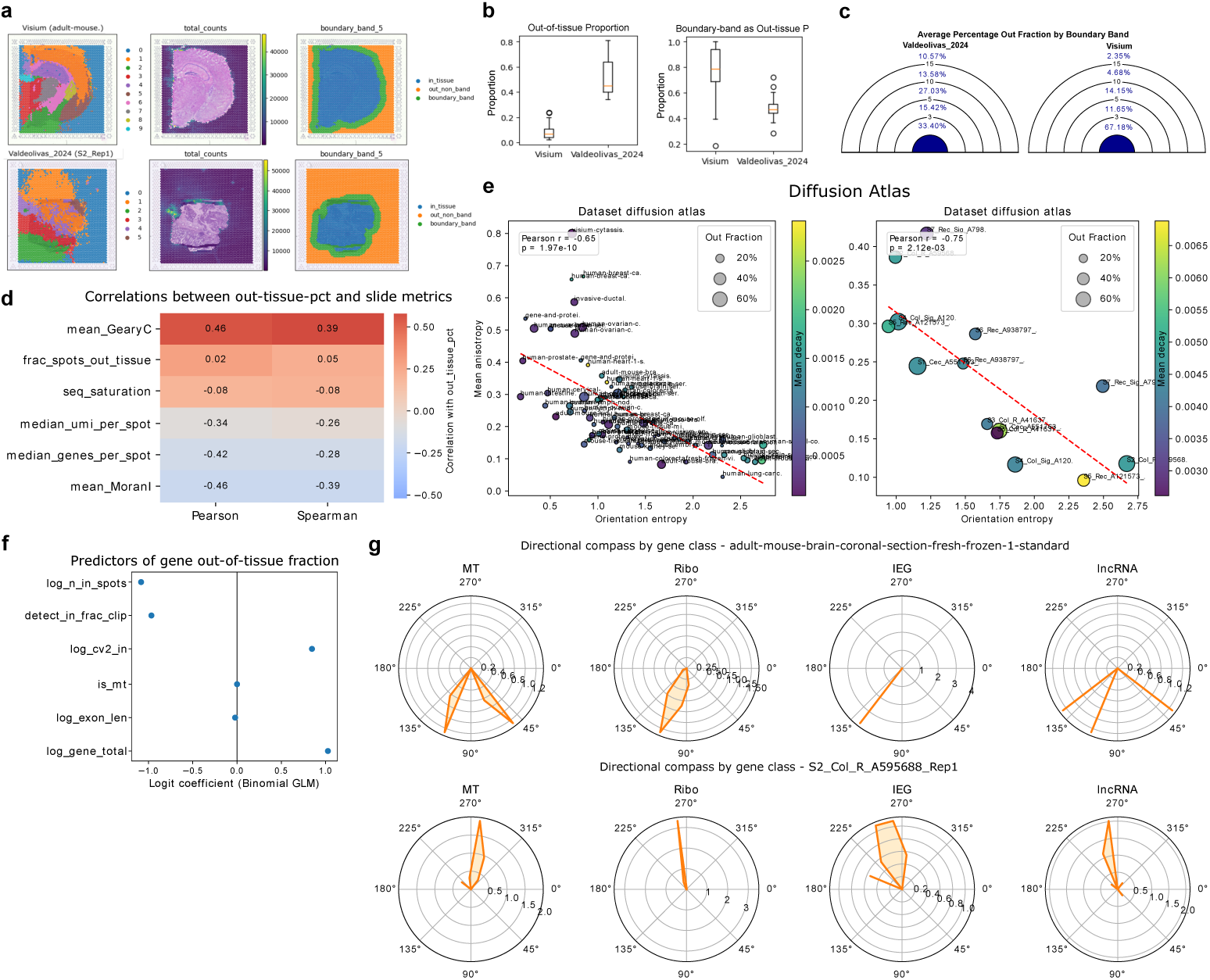
Comparative characterization of mRNA lateral diffusion across 10X and external Visium samples. **a,** Representative slides from each dataset with full-slide unsupervised clustering (left), total-count distribution in out-of-tissue spots (middle), and boundary-band within 5 grid distances (right). Out-of-tissue spots frequently co-cluster with adjacent in-tissue domains, indicating structured rather than random leakage. **b,** Cross-sample distributions of out-of-tissue expression fraction and boundary-band fraction (within 5 grids). Relative to *Valdeolivas 2024*, 10X slides show lower global out-of-tissue burden with higher boundary concentration, consistent with less diffusion with a small, tissue-edge associated halo. **c,** Dartboard-style visualization that partitions out-of-tissue space into concentric bands by multiples of native spot spacing to summarize radial decay from the tissue edge. **d,** Correlation analysis between out-of-tissue expression fraction and sample-level technical/spatial metrics across datasets. Out-of-tissue fraction is weakly related to out-of-tissue spot proportion and sequencing saturation, but negatively associated with median UMI, median detected genes, and Moran’s I, and positively associated with Geary’s C. **e,** Diffusion atlas summarizing sample-level physical diffusion properties. Each point is one slide; x-axis: orientation entropy of preferred diffusion angles (lower entropy indicates more coherent directionality across genes); y-axis: mean anisotropy magnitude; point size: total out-of-tissue fraction; color: mean gene-level distance-decay constant *λ* (higher values indicate steeper decay and more confined leakage). Red dashed line shows linear regression with Pearson *r* and *p*-value annotated. **f,** Binomial generalized linear model of gene-level out-of-tissue fraction with dataset fixed effects, showing covariate coefficients for diffusion propensity (including gene abundance, in-tissue variance-to-mean ratio, detection fraction, number of in-tissue spots, mitochondrial annotation, and exon length). **g,** Directional compass plots of representative gene classes showing preferred diffusion orientations; angular coordinate denotes preferred angle and radial density (the density estimate of the compartment genes at different angles) indicates concentration of genes along each direction.

To summarize diffusion behavior across samples in the two datasets, we first examined the distribution of out-of-tissue expression percentage and boundary band (5 grid distance) count as a fraction of out-of-tissue expression (Fig. 1b). Compared to the *Valdeolivas 2024* dataset, the 10X dataset showed a lower out-of-tissue expression percentage, but a higher boundary band fraction. This demonstrates that the overall diffusion burden is lower in 10X samples, and leaked transcripts are more concentrated near the tissue edge. Although additional independent, external datasets showed similar variability of high out-of-tissue expression, we avoid attributing these differences solely to platform or provider effects, as tissue type, permeabilization conditions and section quality may all influence the apparent out-of-tissue signal, and experimental optimization may require strong expertise. To visualize how out-of-tissue signal varies with distance from the tissue edge, we used a dartboard-style representation (Fig. 1c), which partitions the background region into concentric boundary bands defined in multiples of the native spot spacing. We then calculated correlations between out-of-tissue percentage and some basic parameters and metrics combining both datasets (Fig. 1d). Out-of-tissue expression fraction showed weak association with the fraction of out-of-tissue spots and sequencing saturation. Median UMI, median gene counts and mean Moran’s I showed negative correlations, while the complementary metric Geary’s C showed a positive correlation. These results suggest that diffusion is not simply driven by the number of out-of-tissue spots or sequencing depth. As Moran’s I summarizes spatial autocorrelation, its negative association with out-of-tissue expression fraction is consistent with the possibility that diffusion-like contamination can blur spatially localized expression patterns.

To summarize spatial decay and directional structure in the out-of-tissue signal, we constructed a diffusion-associated atlas across slides (Fig. 1e), in which each point represents one sample slide. The x-axis shows the orientation entropy of preferred diffusion angles across genes within each slide. In contrast to anisotropy, which measures how directional genes are individually, orientation entropy summarizes how similar those preferred directions are across genes: lower entropy indicates that many genes share a coherent diffusion direction, whereas higher entropy indicates that preferred directions are more heterogeneous or closer to isotropic. The y-axis shows the mean anisotropy, which quantifies the average strength of directional bias in out-of-tissue signal across genes: a larger value indicates that, for many genes, the out-of-tissue signal is unevenly distributed along a preferred direction. We observed a significant negative correlation between orientation entropy and mean anisotropy across samples, with the linear regression shown as a red dashed line (Fig. 1e). Point size encodes the overall out-of-tissue fraction and reflects the global burden of diffusion in each sample. Point color represents the mean decay constant *λ* estimated from gene-level distance– decay profiles: warmer colors (higher *λ*) indicate a steeper decline of out-of-tissue signal with increasing distance from the tissue boundary, reflecting more spatially confined background patterns, whereas cooler colors (lower *λ*) indicate broader diffusion halos that extend farther into the background. Thus, samples with high mean anisotropy and low orientation entropy are consistent with directionally coherent out-of-tissue signal. In our dataset, such samples were more frequently observed than samples with both high anisotropy and high orientation entropy. Samples with high anisotropy but higher entropy suggest that directional bias is present but varies across genes, and we can see that this is quite uncommon in both datasets. These observations suggest that, for many slides, gene-level out-of-tissue signals may share a substantial sample-level directional component. This motivated the use of a shared diffusion kernel in CLEAR-ST, while allowing spot-specific diffusion rates to capture local heterogeneity.

To identify gene-level properties associated with diffusion, we next modeled the gene out-of-tissue fraction using a binomial generalized linear model with dataset fixed effects (Fig. 1f). We observed that total gene counts (log gene total) and the in-tissue variance-to-mean ratio (log cv2 in) were among the strongest positive predictors of gene-level out-of-tissue fraction (Fig. 1f). The former is expected, as highly abundant genes have more transcripts available to diffuse, whereas the latter suggests that genes with more heterogeneous expression across in-tissue spots are more likely to exhibit larger out-of-tissue fractions. One possible interpretation is that spatially heterogeneous in-tissue expression creates stronger local gradients, which could increase the probability of observing boundary-associated spillover. By contrast, both the in-tissue detection fraction (detect in frac clip) and the number of in-tissue spots in a dataset (log n in spots) showed strong negative effects. We interpret these as structural rather than molecular determinants: genes detected broadly across many in-tissue spots have a larger in-tissue denominator, so a similar amount of leakage contributes a smaller fraction of their total counts, while datasets with more in-tissue spots likely contain a larger tissue interior and are therefore less dominated by boundary-associated spillover. Lastly, mitochondrial identity (is mt) and exon length (log exon len) had coefficients close to zero in this model, suggesting that these covariates did not explain substantial variation in gene-level out-of-tissue fraction after accounting for the other included predictors.

Finally, to examine whether diffusion is directionally biased, we summarized representative gene-class-specific preferred diffusion directions using directional compass plots (Fig. 1g). In these polar plots, the angular coordinate denotes the preferred diffusion direction of a gene, estimated from the orientation of out-of-tissue signal relative to the nearest tissue boundary, and the radial profile reflects the density of genes with preferred angles in that direction. A sharp peak therefore indicates that genes within a class tend to show out-of-tissue signal enriched along a common direction, whereas a flatter or more uniform profile indicates either weak directionality or greater heterogeneity in diffusion orientation. These compass plots were visually consistent with the spatial distribution of expression-derived clusters and total counts in representative slides (Fig. 1a). For example, in the S2 Rep1 sample, the estimated preferred direction was oriented toward the positive *y*-axis (Fig. 1g, bottom), and the spatial pattern of cluster 1 is consistent with this directional bias: the out-of-tissue spots co-clustered with cluster 1 are mostly located above the tissue boundary, and lower right spots adjacent to cluster 1 show weak co-clustering (Fig. 1a, bottom left).

Overall, these analyses indicate that out-of-tissue expression is frequently spatially structured, varies substantially across datasets, and can exhibit directional coherence within individual samples. These observations motivated three design choices in CLEAR-ST: expression-derived initialization of spot-specific diffusion rates, histology-derived niche labels as auxiliary structural information, and a shared graph-Laplacian-based diffusion kernel to capture sample-level spatial spread while retaining spot-level heterogeneity.

### 2.2 CLEAR-ST models diffusion-like contamination using a denoising autoencoder coupled with a forward diffusion framework

Motivated by the structured and often directionally coherent out-of-tissue signal described above, CLEAR-ST is designed to infer an underlying expression field whose forward diffusion can reproduce the observed contaminated counts. CLEAR-ST takes as input the raw spatial transcriptomics count matrix, spot coordinates, the corresponding histology image, and a tissue mask (Fig. 2b). These inputs are routinely available for Visium-like capture-based spatial transcriptomics datasets.

**Fig. 2.**
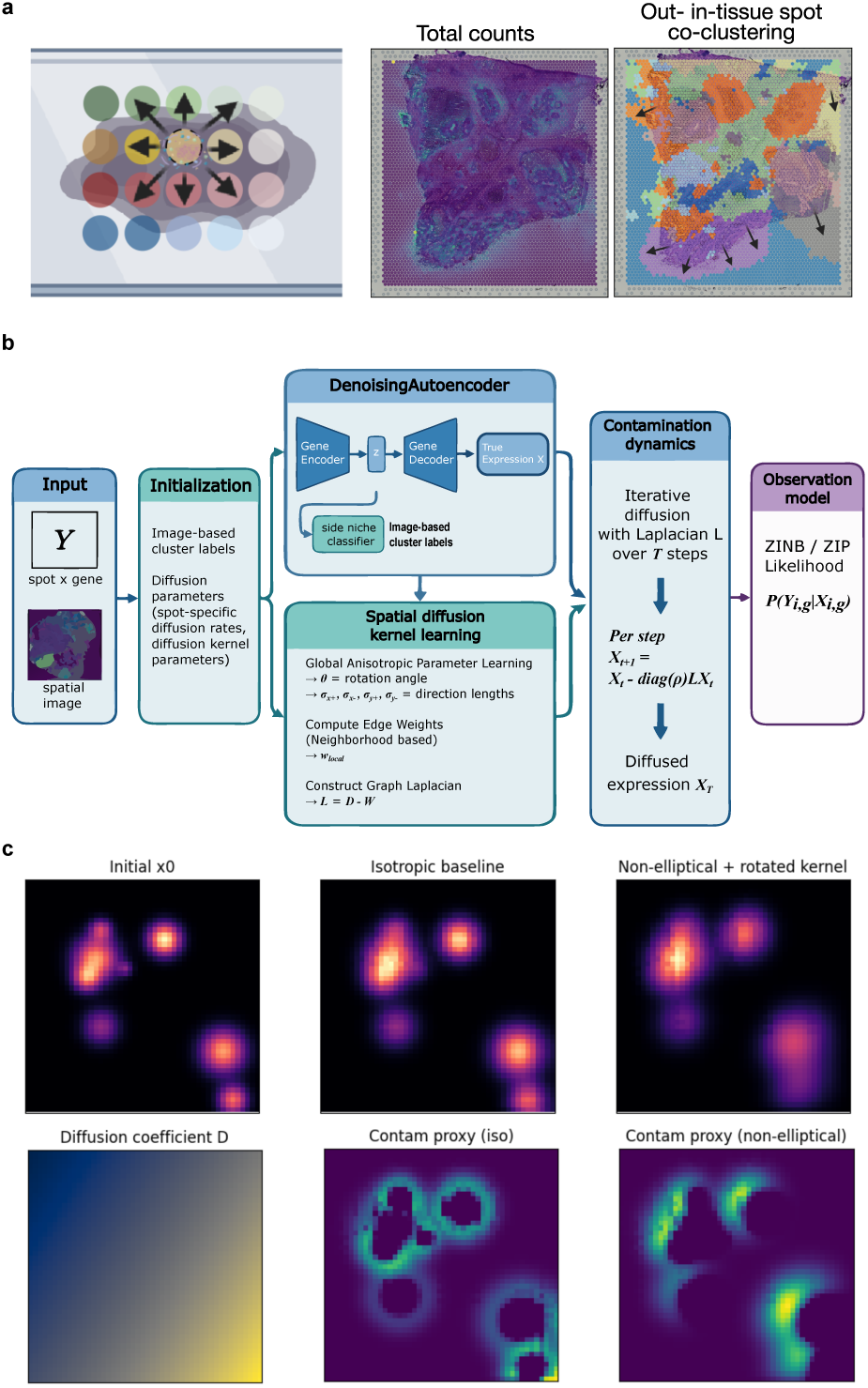
Overview of the CLEAR-ST model. **a,** Illustration of the mRNA lateral diffusion issue. During the sequencing process the tissue slice is permeabilized and mRNA molecules are allowed to diffuse in the solution. After data processing, we can detect a certain percentage of mRNA counts located in out-of-tissue spots, especially those adjacent to in-tissue spots. Clustering of out- and in-tissue spots together often presents as this “spilling-out” pattern as out-of-tissue spots appears transcriptomically similar to adjacent in-tissue spots. **b,** Model design of the CLEAR-ST model. It mainly comprised a denoising autoencoder and a forward contamination process to derive contaminated counts from imputed clean expression. **c,** Demonstration of using anisotropic Laplacian with spatially-varying diffusion coefficients in forward diffusion with a simulated field with scattered substance mass. Top row shows the initial field, isotropic diffusion endpoint distribution, and anisotropic diffusion endpoint distribution. The bottom row shows the mass diffused-out for isotropic and anisotropic scenarios, and the diffusion coefficients which we set to be increasing with *x* and *y* coordinates.

During preprocessing, the histology image is cropped into spot-level patches, which are clustered to obtain image-derived niche labels. These labels are intended to provide auxiliary information about local tissue architecture that is independent of transcript capture artifacts, although they may still be affected by staining quality, tissue morphology, or image-processing errors. In parallel, expression profiles are clustered to derive expression-based initialization labels, allowing out-of-tissue spots to be associated with transcriptionally similar in-tissue regions when such structure is present (Fig. 2b). These labels are used only to initialize spot-specific diffusion-rate parameters, which are subsequently updated during probabilistic inference. A user-defined number of highly variable genes is selected for model fitting to reduce dimensionality and focus inference on genes with informative spatial and transcriptional variation.

The core model then couples a denoising autoencoder with a forward diffusion framework: the autoencoder encodes the observed gene expression matrix and decodes an estimated clean-expression field. The forward diffusion module then applies a graph-Laplacian-based operator to this estimated clean-expression field, using spot-specific diffusion rates and learnable kernel parameters to model spatial spread consistent with diffusion-like contamination (Fig. 2c). Physically, this forward formulation reflects the actual experimental sequence: transcripts first reside at their true cellular locations (the clean expression field), then spread outward through the tissue during permeabilization, and are finally captured at probe-coated spots. We model this forward process - transcripts moving from clean sources to contaminated observations - in a physically faithful way, simulating the observed spot counts which integrate contributions from multiple source locations. Within this framework, each entry *w_ij_* of the graph weight matrix **W** represents the fraction of transcripts originating at spot *i* that diffuse to neighboring spot *j* in a single step, with the row-stochastic normalization *Σ_j_w_ij_ = 1* ensuring conservation of total transcript mass during spatial redistribution. The resulting contaminated-expression estimate is linked to the observed counts through an explicit probabilistic observation model, with a Zero-inflated Poisson likelihood used by default and alternative count likelihoods, such as Zero-inflated Negative Binomial, available as options. We use the ZIP likelihood as the default for computational stability, while allowing users to select more flexible likelihoods when overdispersion is substantial. During training, all model components are optimized jointly by stochastic variational inference to maximize the probabilistic agreement between the observed counts and the counts predicted after the forward contamination process, so that the autoencoder estimates expression patterns that, after the modeled contamination process, best explain the observed data. This joint optimization yields estimated decontaminated expression profiles, inferred diffusion-related parameters, and latent tissue representations.

### 2.3 CLEAR-ST improves spatial structure and domain identification in diffusion-affected data

We evaluated CLEAR-ST on samples from the *Valdeolivas 2024* dataset, which exhibits more severe diffusion than standard 10x Visium slides. In parallel, we applied SpotClean and resolVI as the benchmarked methods.

For a representative slide (S6 Rec A938797 Rep1; 34% out-of-tissue expression), we visualized the raw total UMI counts (Fig. 3b) together with the inferred spot diffusion-rate groups (Fig. 3c). The high-diffusion group was mainly comprised of tumor and stroma fibroblastic IC high regions (Fig. 3c). Checking the directional bias, we found large sigmas in X axis positive direction and Y axis negative direction, while the other two direction arms (X negative and Y positive) had comparatively smaller sigmas (35.30, 27.56 compared to 1.45 and 2.92). These parameter values approximately aligned with the observed total count patterns on the slide (Fig. 3b), suggesting that the forward diffusion process of CLEAR-ST learned the anisotropic diffusion pattern during optimization. We then performed unsupervised Leiden clustering on both the raw and CLEAR-ST-corrected expression matrices across a range of resolutions (0.1–1.5, with 0.05 as the step size), and compared the resulting partitions using silhouette scores and annotation-based clustering metrics relative to the pathologist labels (Fig. 3a), while keeping the number of clusters matched across methods (Fig. 3d,e). Across cluster numbers, CLEAR-ST-corrected expression generally yielded higher silhouette scores, indicating increased within-cluster similarity and greater separation between clusters in the embedding space (Fig. 3d). Concordantly, we evaluated clustering results based on pathologist annotations using normalized mutual information (NMI), adjusted rand index (ARI) and cluster Homogeneity (Fig. 3e). We observed a consistent improvement after CLEAR-ST correction, suggesting improved agreement with the annotated spatial domains identified by expert histopathological assessment. SpotClean and resolVI showed more modest to none improvements in clustering metrics, with resolVI unable to produce cluster number lower than 8. For the best annotation-matching resolution (*k* = 6), we visualized both the spatial cluster assignments and spot-level silhouette scores (Fig. 3f,g). CLEAR-ST increased silhouette scores particularly in spots located within the stroma fibroblastic IC and stroma desmoplastic IC low regions, consistent with improved delineation of these domains after correction. Before correction, stroma fibroblastic IC was split across clusters 1 and 2, whereas stroma desmoplastic IC low was not recovered as a distinct spatial cluster (Fig. 3f). After correction, these two domains were more clearly resolved as clusters 0 and 5, respectively (Fig. 3g).

**Fig. 3.**
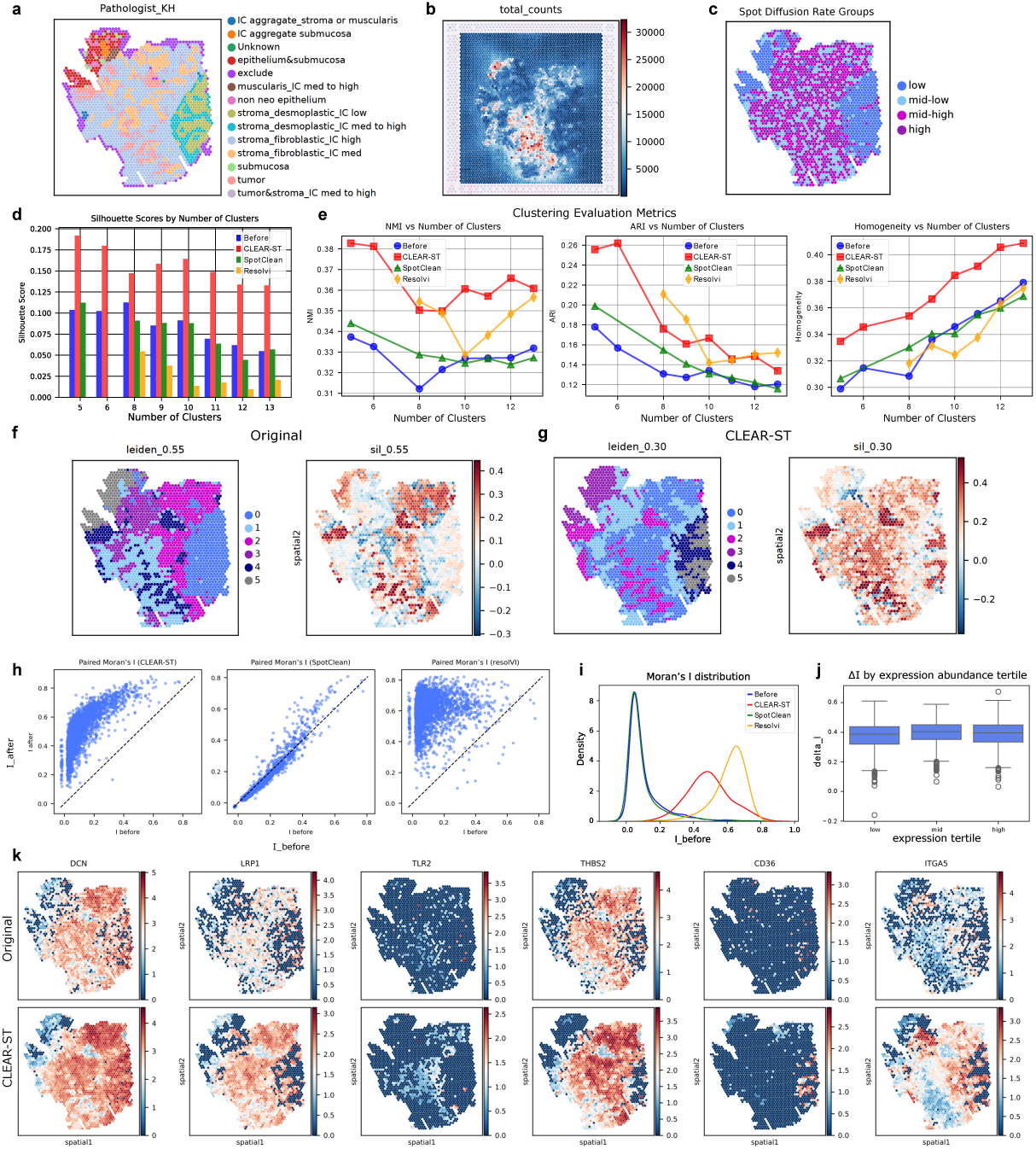
CLEAR-ST correction improves clustering quality and spatial structure recovery in a colorectal cancer slide (S6 Rec A938797 Rep1; 34% out-of-tissue expression), compared with SpotClean and resolVI. **a,** Pathologist annotations of the sample. **b,** Raw total UMI distribution across the slide. **c,** Inferred spot diffusion-rate groups, showing spatial concordance with high-count regions and tissue boundary diffusion context. **d,** Silhouette score profiles across cluster numbers before and after CLEAR-ST correction, and for clustering results based on SpotClean and resolVI corrected expression. **e,** Clustering metrics across methods at matched cluster-number settings (CLEAR-ST, SpotClean, resolVI), including NMI, ARI and Homogeneity. **f,** Spatial cluster assignments for leiden resolution 0.55 before correction (*k* = 6) with spot-level silhouette scores. **g,** Spatial cluster assignments for leiden resolution 0.30 after correction (*k* = 6) with spot-level silhouette scores. **h,** Gene-level Moran’s I distributions before versus after correction for each method (CLEAR-ST, SpotClean, resolVI). **i,** Pergene Moran’s I change summary (Δ*I*) highlighting broader positive shifts for CLEAR-ST (and similarly for resolVI) relative to SpotClean. **j,** Stratified Δ*I* by expression quantile, showing improvements that are not restricted to highly expressed genes. **k,** Representative tumor-related spatial expression maps before and after correction, illustrating sharper localization and reduced diffusion blur after CLEAR-ST correction.

We next examined the distribution of Moran’s I values before and after correction, motivated by our earlier observation that stronger diffusion is associated with reduced spatial autocorrelation (Fig. 3h,i). CLEAR-ST led to a shift toward higher Moran’s I values across genes (Fig. 3h, left). resolVI produced a qualitatively similar increase, whereas SpotClean resulted in comparatively little change in the Moran’s I distribution (Fig. 3h, middle and right). These results suggest that CLEAR-ST and resolVI are more effective at restoring spatial structure blurred by diffusion, while SpotClean has a more limited impact on gene-level spatial organization. To assess whether these improvements depended on expression abundance, we further stratified changes in Moran’s I by expression quantile (Fig. 3j). We observed no clear dependence of Moran’s I improvement on expression level, indicating that CLEAR-ST does not preferentially enhance highly expressed genes but instead improves spatial structure across the expression spectrum. Finally, we examined the spatial patterns of representative tumor-related genes before and after correction (Fig. 3k). Following CLEAR-ST correction, these genes displayed more spatially localized expression, consistent with the observed increase in Moran’s I and with the improved identification of the stroma fibroblastic IC and stroma desmoplastic IC low regions. We also visualized genes with the highest improvement in Moran’s I, which showed a much clearer spatial pattern after correction (Supplementary Fig. 1a,b). For instance, PTEN11’s localization became much better aligned with the tumor region, which makes biological sense given its known role as a tumor suppressor gene [13].

Overall, these results suggest that CLEAR-ST can improve spatial structure and domain separability, leading to more accurate gene expression profiles that enhance the identification of biologically relevant spatial domains, aiding bioinformatic analysis and facilitating easier interpretation of spatial transcriptomics data.

### 2.4 Differential expression and gene set enrichment analyses support the biological interpretability of CLEAR-ST-corrected expression profiles

To further assess whether CLEAR-ST-corrected expression profiles preserve and clarify biologically interpretable signals, we performed downstream marker-gene and pathway analyses using corrected counts. We applied CLEAR-ST to an additional colorectal cancer slide from the same dataset (S4 Col Sig A120838 Rep2; 65% out-of-tissue expression; Fig. 4a). The high-diffusion group is mainly comprised of stroma fibroblastic IC high and stroma fibroblastic IC med regions (Fig. 4a). In terms of diffusion kernel sizes, X axis’ negative arm showed the highest sigma value (12.61), while sigma values of the other three arms were in the range of 5–9, which aligned with the observed total counts and co-clustering labels (Supplementary Fig. 2a). We examined gene-level Moran’s I distributions and silhouette scores across cluster resolutions, which showed trends similar to those observed in the previous slide (Supplementary Fig. 2a–c). Relative to the evaluated benchmark methods, CLEAR-ST showed more consistent increases in annotation-alignment metrics, including ARI, NMI, and Homogeneity, in this slide (Supplementary Fig. 2b). The largest apparent gains in silhouette score and annotation agreement were observed around the stroma fibroblastic IC high region, defined as a fibroblast-rich, high immune infiltration and tumor-stroma interface region by the original paper [12]. Before correction, cluster 1 overlapped with both the low-confidence region labelled “exclude” and part of the tumor region (Fig. 4b), and this mixing was still observed at higher clustering resolution (Supplementary Fig. 2c). The tumor region also showed higher silhouette scores after correction, consistent with increased within-region similarity and greater separation from other expression-defined regions.

**Fig. 4.**
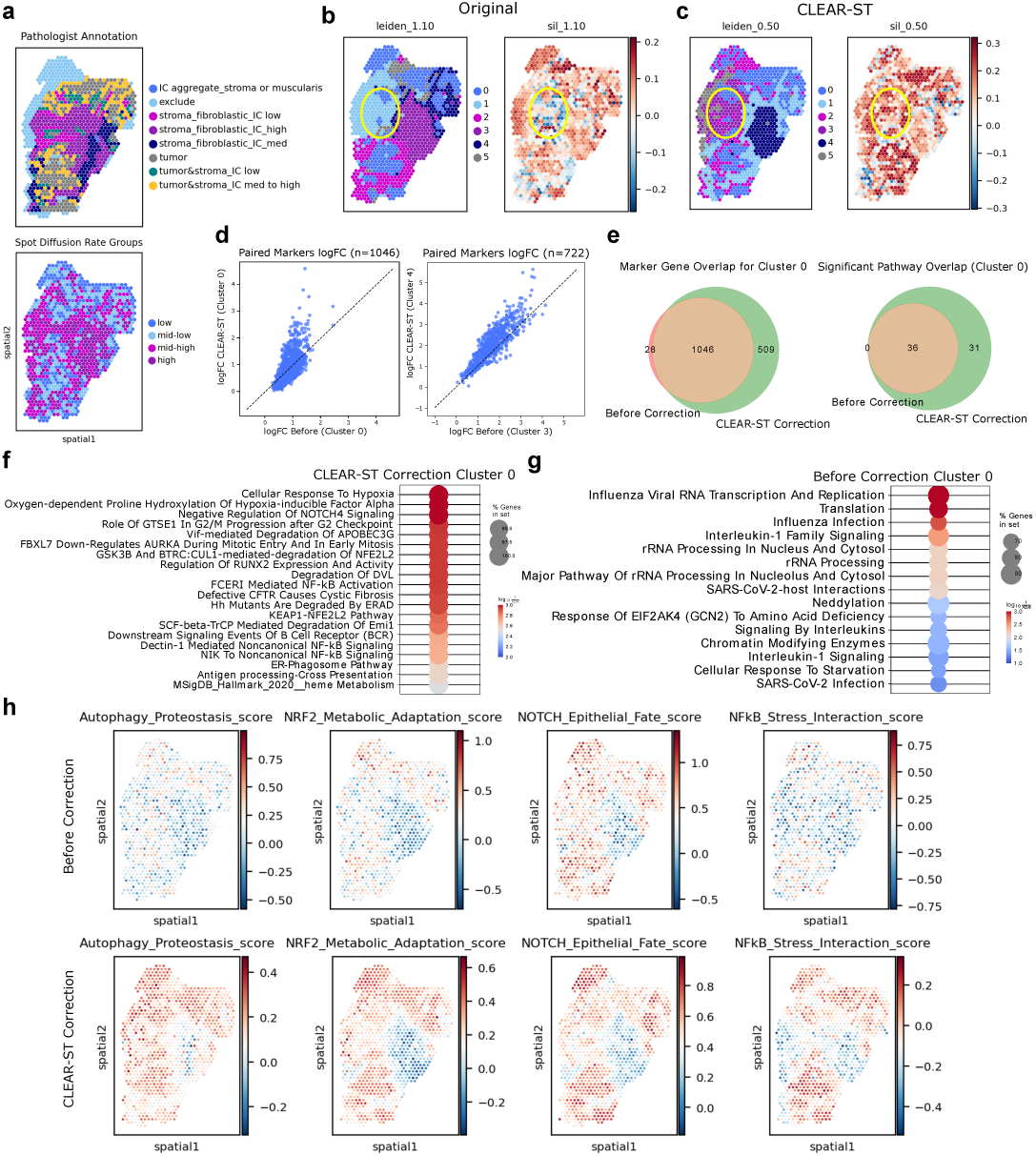
Downstream marker and pathway analyses on CLEAR-ST-corrected expression reveal improved biological specificity in a high-diffusion colorectal slide (S4 Col Sig A120838 Rep2). **a,** Pathologist annotations and the histological image of the sample. **b,** Spatial clustering maps before correction, highlighting incomplete separation of stromal and tumor-adjacent compartments. **c,** Spatial clustering maps after correction, showing clearer separation of biologically interpretable domains, including stromal subregions. **d,** Matched-cluster marker effect-size (logfoldchanges) comparison (before vs after), where each point is a shared marker and axes denote log fold-change estimates from the two conditions. **e,** Marker and pathway overlap summaries (Venn-style) for representative matched clusters/tumor compartment, demonstrating retention of core signals with additional discoveries after correction. **f,** Gene sets significantly enriched only after correction (FDR *<* 0.05). **g,** Shared significantly enriched pathways before and after correction, mainly core tumor biosynthetic programs. **h,** Spatial module-score maps for representative pathway modules (top: raw; bottom: corrected); corrected profiles show reduced diffuse extremes and sharper niche-aligned spatial structure.

To evaluate and compare markers discovered under the before and after correction scenarios, we matched raw and CLEAR-ST-corrected clusters using a spot-overlap criterion, pairing clusters with more than 80% overlap. We found cluster 3 (raw) – cluster 4 (corrected) and cluster 0 (raw) – cluster 0 (corrected) as matching clusters, and compared the paired logfoldchanges before (*x* axis) and after correction (*y* axis) (Fig. 4d). The matched cluster pair cluster 3 (raw)–cluster 4 (corrected) corresponded primarily to the stroma fibroblastic region. This pair contained 728 significant markers before correction and 1126 after correction, with 722 markers shared between the two analyses. Among shared markers, the average log-fold-change increased by 0.343 after correction. Similarly, the matched tumor-associated pair cluster 0 (raw)–cluster 0 (corrected) contained 1074 significant markers before correction and 1555 after correction, with 1046 shared markers and a smaller average increase in shared-marker log-fold-change of 0.029 (Fig. 4e). We therefore interpreted increases in marker number together with marker overlap and effect-size changes, rather than treating the number of significant markers alone as evidence of improved biological specificity.

We next compared significantly enriched gene sets in the matched tumor-associated cluster before and after correction to assess whether pathway-level signals were preserved or clarified after correction. In this matched comparison, all gene sets significant in the raw analysis remained significant after correction, while 31 additional gene sets reached significance only in the corrected analysis (Fig. 4e). Gene sets shared between before and after correction were associated with core cellular machinery, including mRNA translation, ribosomal subunit assembly, translation initiation and elongation, nonsense-mediated decay, cotranslational protein targeting, and cytokine signaling (Fig. 4g). The retention of these fundamental processes suggests that correction did not remove major tumor-associated biosynthetic and signaling programs in this comparison. In contrast to the shared pathways, gene sets that reached significance only after correction included several biologically interpretable tumor-associated programs. These included autophagy and macroautophagy, NRF2-mediated oxidative stress responses, hypoxia and heme metabolism pathways, and regulatory signaling through NOTCH and WNT-*β*-catenin axes (Fig. 4f). In addition, pathways related to controlled cell-cycle progression, DNA damage checkpoints, ubiquitin-mediated proteolysis, and p53 stabilization became evident only after correction (Fig. 4f). These pathways are consistent with processes commonly implicated in colorectal cancer biology, including metabolic stress responses, redox regulation, epithelial fate control, and cell-cycle regulation. Importantly, the appearance of these additional pathways was accompanied by retention of core biosynthetic and signaling pathways, arguing against a wholesale loss of major tumor-associated programs in this analysis. These results suggest that correction may improve the detectability of pathway-level signals that are less apparent in the raw data, while reducing the influence of diffuse background expression. To better visualize the impact of CLEAR-ST on these gene modules, we calculated module scores with each gene set and visualized their spatial patterns (Fig. 4h, top – raw, bottom – corrected). Before correction, module-score maps appeared spatially diffuse and showed limited visual correspondence with annotated spatial niches (Fig. 4h, top). After correction, extreme module-score values were reduced, and the resulting spatial patterns showed stronger visual correspondence with annotated niches (Fig. 4h, bottom).

In conclusion, these results show that CLEAR-ST enhances the resolution and interpretability of gene set enrichment analyses by preserving essential tumor cell traits while uncovering additional layers of tumor-intrinsic adaptation and regulatory control that are obscured in raw expression profiles.

### 2.5 CLEAR-ST-corrected expression profiles improve the spatial coherence of cell type deconvolution

To assess whether CLEAR-ST correction enhances the biological interpretability of inferred cell type organization, we further evaluated whether corrected spatial counts align better with single-cell references and yield cell type distributions that are more consistent with known spatial organization of thymic cell populations. To diversify sample sources, we applied CLEAR-ST to another public dataset, and present here the result of one slide (sample name: TA11486163, 13% out-of-tissue expression) [14]. The fitted diffusion rates of this slide were relatively uniform and the diffusion rate groups were spatially intermixed (Supplementary Fig. 3a). The diffusion kernel widths were approximately 30 for all arms, indicating an isotropic diffusion behavior on this slide. Despite the relatively low out-of-tissue expression percentage, we observed modest improvements in the spatial delineation of thymic niches by cluster matching metrics NMI and ARI (Supplementary Fig. 3c). Specifically, after correction, cluster 0 more clearly corresponded to medullary-associated regions enriched for mature immune populations, while cluster 1 aligned more closely with capsular or peripheral stromal compartments, accompanied by silhouette score improvements in these regions (Supplementary Fig. 3a,b).

We quantified this niche-identification improvement in two ways with information from deconvoluted cell abundance. First, for each spot we assigned the dominant inferred cell type (arg max*_c_ x_i,c_*, where *x_i,c_* is the abundance of cell type *c* in spot *i*) and measured agreement with pathologist regions using normalized mutual information (NMI), which increased from 0.162 to 0.175 (Δ = 0.013). Second, we trained a logistic-regression classifier to predict pathologist labels from cell-type abundance and evaluated performance by stratified 5-fold cross-validation (i.e., five train/test splits preserving region proportions in each split). Both metrics showed modest increases after correction, consistent with improved alignment between inferred cell type distributions and annotated anatomical regions (mean accuracy: 0.262 → 0.284; macro-F1: 0.245 → 0.263).

From a technical perspective, we first inspected posterior uncertainty by quantifying the posterior width of inferred cell type abundances. For each spatial location and cell type, posterior width was computed as the difference between the 95th and 5th percentile of the posterior distribution estimated by Cell2location (Fig. 5b). For each metric below, before-versus-after differences were tested by paired Wilcoxon signed-rank tests, with Benjamini–Hochberg false-discovery-rate correction across related tests. Posterior uncertainty decreased across spots, indicating more stable inference of dominant cell type contributions (*n* = 2831): median posterior mean width (median*_c_{µ_i,c_*}, where *µ_i,c_* is the mean width across cell types) decreased from 0.599 to 0.488 (FDR *<* 1 × 10*^−^*^300^) (Fig. 5b). We next evaluated the dominance of the most probable cell type per spot using the top-1 posterior probability, defined as the maximum normalized cell type abundance at each spatial location (Fig. 5c). Higher top-1 probabilities reflect reduced ambiguity in assignment and greater confidence that a single cell type predominantly explains the observed expression profile at that spot. Compared to raw counts, corrected expression profiles yielded systematically higher top-1 probabilities (median 0.0586 → 0.0884, FDR *<* 1 × 10*^−^*^300^), suggesting that individual spots were more strongly dominated by a limited number of cell types, consistent with the expected spatial organization of thymic microenvironments (Fig. 5c). To capture uncertainty in a distribution-aware manner, we further computed Shannon entropy of the normalized posterior cell type proportions for each spot which provides a continuous measure of how evenly probability mass is distributed across cell types, with higher values indicating greater ambiguity (Fig. 5d). After correction, entropy values decreased across the majority of spots (median normalized entropy 0.966 → 0.935, FDR *<* 1 × 10*^−^*^300^), consistent with a sharpening of cell type assignments and a reduction in spurious cell type mixing introduced by mRNA diffusion. Finally, to assess whether improved deconvolution confidence translated into more spatially coherent cell type organization, we computed Moran’s I for each inferred cell type abundance before and after correction, with higher values suggesting more localized and meaningful spatial patterns. Again, we saw that most cell types showed an increase in Moran’s I after correction: 54 of 56 inferred cell types had positive Δ*I* (median Δ*I* = 0.062; mean Δ*I* = 0.066), with largest gains in pDC (Δ*I* = 0.188), TEC-myo (Δ*I* = 0.165), and aDC3 (Δ*I* = 0.163). Only T CD4 and T CD8 showed decreases (Δ*I* = 0.099 and 0.127, respectively), indicating that the inferred spatial distribution of most cell types became more spatially coherent (Fig. 5e). Overall, these metrics indicate that CLEAR-ST-corrected expression profiles not only improve posterior certainty but also enhance the recovery of spatially structured cell type distributions.

**Fig. 5.**
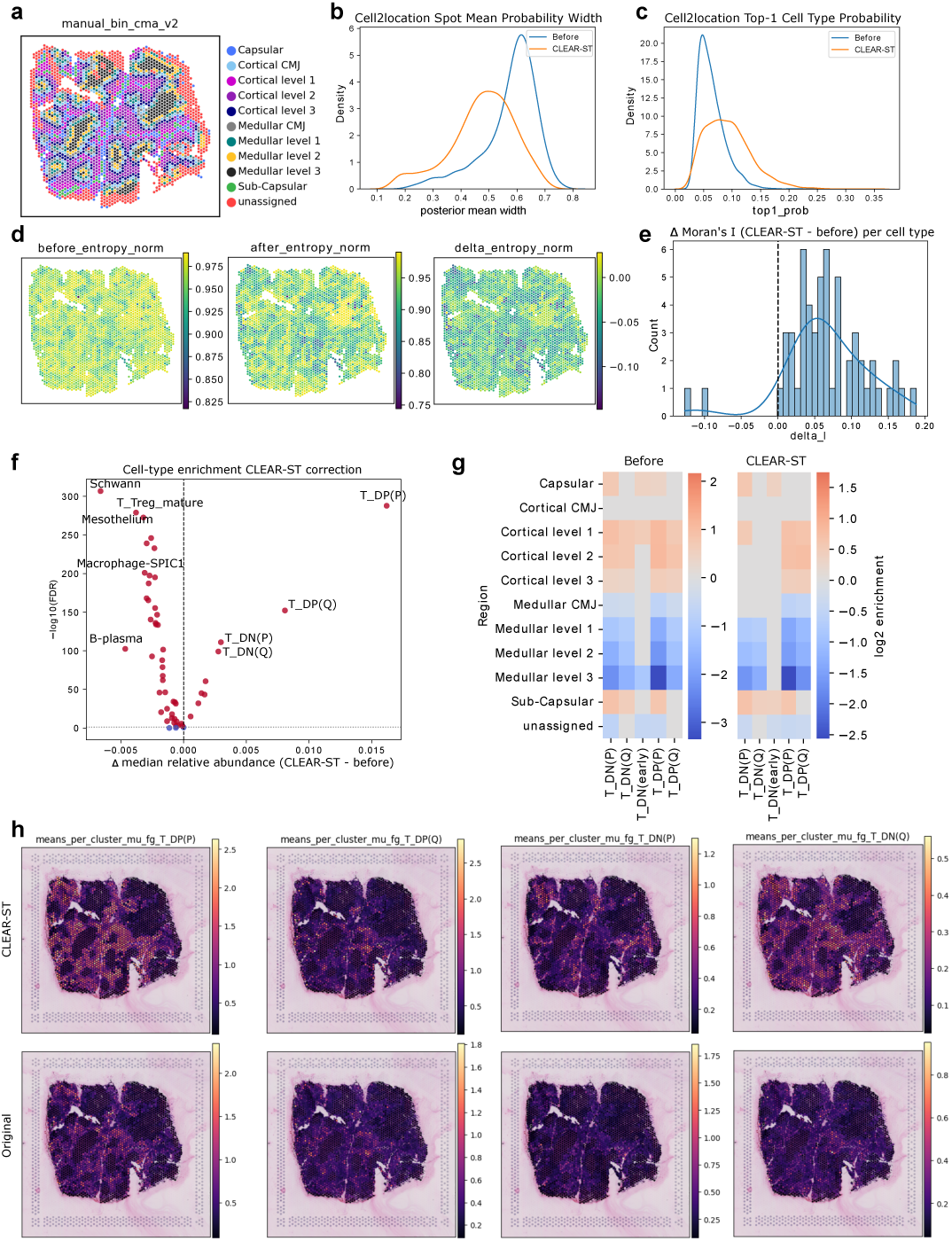
Cell2location evaluation demonstrates improved confidence, spatial coherence, and biological interpretability after CLEAR-ST correction. **a,** Manual annotations of thymic compartments from the original study. **b,** Posterior uncertainty of inferred cell-type abundance, quantified as posterior width (*q*_95_ *− q*_05_) means for each spot across cell type. CLEAR-ST reduces uncertainty across spots (median posterior mean width 0.599 *→* 0.488). **c,** Top-1 posterior dominance per spot (maximum normalized inferred abundance across cell types), showing increased assignment confidence after correction (median 0.0586 *→* 0.0884). **d,** Shannon entropy of normalized posterior cell-type proportions per spot, indicating reduced ambiguity after correction (median normalized entropy 0.966 *→* 0.935). **e,** Change in Moran’s I of inferred cell-type abundance maps before versus after correction, showing improved spatial autocorrelation for most cell types (54/56 with Δ*I >* 0; median Δ*I* = 0.062). **f,** Volcano plot of changes in median relative cell-type abundance after correction; points are annotated by effect size and FDR-significance (paired Wilcoxon, Benjamini–Hochberg adjusted), highlighting increased T DN/DP thymocyte programs and depletion of diffuse background populations. **g,** Region-by-cell-type niche enrichment heatmaps before and after correction. **h,** Spatial maps of representative T-cell subtype abundances before and after correction. After correction, T DN populations show increased relative enrichment in subcapsular regions with reduced apparent spread into cortical areas, while T DP populations remain predominantly distributed across cortical compartments.

We then investigated which cell types became more or less abundant after correction by visualizing changes in median relative abundance using a volcano plot (Fig. 5f). Of 56 inferred cell types, 53 showed significant shifts in median relative abundance (paired Wilcoxon with Benjamini–Hochberg correction, FDR *<* 0.05): 9 increased and 44 decreased. The largest increases were observed in T DP(P) (Δ median relative abundance = 0.0162, FDR = 1.59 × 10*^−^*^288^), T DP(Q) (Δ = 0.00807, FDR = 5.93 × 10*^−^*^153^), T DN(P) (Δ = 0.00297, FDR = 8.15 × 10*^−^*^112^), and T DN(Q) (Δ = 0.00277, FDR = 7.85 × 10*^−^*^100^), all of which represent distinct stages of developing thymocytes. These populations correspond to well-characterized phases of early T-cell maturation, progressing from proliferative and quiescent double-negative (DN) stages that localize predominantly to the subcapsular region, to proliferative and quiescent double-positive (DP) stages that are distributed across the cortical compartment of the thymus prior to selection [14]. At the whole-slide level, the average composition shift between before and after was moderate (Jensen–Shannon distance = 0.082). To assess how spatial associations changed after correction, we compared niche enrichment profiles before and after correction using a heatmap (Fig. 5g), with the full version shown in Supplementary Fig. 3f. Niche enrichment was quantified using a log_2_ enrichment score, defined as the base-2 logarithm of the ratio between the mean abundance of a given cell type within a spatial region and its mean abundance outside that region. We visualized the top 15 cell types ranked by enrichment score in each thymus region. Using this metric, we observed that after correction, T_DN(P) showed increased relative enrichment in the subcapsular zone, accompanied by a reduction in apparent enrichment across cortical regions. This pattern is consistent with the known biology of DN thymocytes, which preferentially localize to subcapsular niches while remaining spatially distributed due to ongoing migration. T_DN(Q) and early DN thymocytes (T_DN(early)) similarly showed predominant enrichment in subcapsular regions, with reduced representation among the most enriched populations in cortical compartments after correction (Fig. 5g). Both DP populations, T_DP(P) and T_DP(Q), remained enriched across cortical levels 1–3, consistent with their established cortical distribution prior to positive selection. Notably, these cortical enrichment patterns were preserved after correction, suggesting that CLEAR-ST primarily refines the separation between DN and DP spatial profiles rather than altering the dominant localization of DP populations. In parallel, several populations were significantly depleted, including neutrophils (Δ = −0.0105, FDR *<* 1 × 10*^−^*^300^), Schwann cells (Δ = −0.00663, FDR = 1.50 × 10*^−^*^307^), B-plasma (Δ = −0.00466, FDR = 3.10 × 10*^−^*^103^), and mature Treg (Δ = −0.00380, FDR = 1.04 × 10*^−^*^279^), suggesting reduced diffuse background contributions from non-dominant compartments. Finally, visualization of the spatial distributions of these T-cell subtype abundances showed that, following correction, their spatial patterns became more localized and consistent with established thymic biology, indicating improved correspondence between inferred cell-type localization and known developmental niches (Fig. 5h). We also conducted the same investigation for the previous sample S4 Col Sig A120838 Rep2 (Supplementary Fig. 2d–g). Posterior related metrics, cell type entropy, and cell type Moran’s I were shown to be improved in a similar fashion. The most enhanced cell type was stem cell, with the most increased enrichment observed in stroma fibroblastic IC high and tumor regions (Supplementary Fig. 2f,g).

Overall, these results suggest that CLEAR-ST-corrected expression profiles are associated with more spatially coherent and biologically consistent cell type deconvolution outputs, with inferred cell type distributions showing improved correspondence to known thymic tissue organization.

## 3 Discussion

In this study, we presented CLEAR-ST as a physics-informed probabilistic decontamination framework for spatial transcriptomics and showed that it improves downstream biological analysis across multiple datasets and analysis tasks. We first performed a detailed analysis regarding the physical diffusion patterns, revealing that unsymmetric diffusion patterns can take place in mRNA lateral diffusion (high anisotrophy), and that genes generally diffuse under the same physical condition (low orientation entropy). These observations motivated the design of CLEAR-ST and we provide these visualization tools as part of CLEAR-ST to help researchers understand the mRNA diffusion patterns in their custom datasets.

Across slides with distinct diffusion burdens, CLEAR-ST consistently increased clustering quality (silhouette and annotation-alignment metrics), shifted gene-level Moran’s I toward higher values, improved marker recovery and effect-size stability, and yielded more interpretable GSEA outputs. Importantly, improvements were not limited to one endpoint: they propagated from low-level spatial structure to high-level biological interpretation and cell-type deconvolution.

Compared with SpotClean and resolVI, CLEAR-ST showed a favorable balance between restoring spatial structure and preserving biologically meaningful signal. SpotClean provided a useful baseline for count redistribution, but in our evaluation its impact on Moran’s I and clustering consistency was generally more modest. resolVI frequently improved spatial autocorrelation and latent representation quality, but CLEAR-ST showed more stable behavior across cluster resolutions and stronger gains in annotation concordance in the tested slides. These differences are consistent with CLEAR-ST’s design: by coupling autoencoding with an explicit graph-based forward diffusion operator, the model can jointly represent expression denoising and physically constrained transcript spread rather than relying on only one mechanism.

The Cell2location analyses further support this conclusion. Using CLEAR-ST-corrected inputs, posterior abundance estimates became sharper, with lower entropy, larger top-1 dominance, narrower posterior quantile widths, and improved spatial coherence of inferred cell-type maps. Region-level enrichment patterns and region-classification performance were also more aligned with known tissue biology. Together with the marker/GSEA findings, these results suggest that CLEAR-ST does not merely smooth expression, but can improve signal specificity in a way that enhances both statistical confidence and biological interpretability.

Despite these strengths, CLEAR-ST has several limitations. First, it relies on image-derived niche labels as an auxiliary prior. If histology quality is poor, if staining artifacts are present, or if image-based clustering does not reflect transcriptional microenvironments, this prior may bias optimization and reduce correction quality in some regions (for which we recommend users to lower image niche regularization strengths). Second, diffusion-rate initialization depends on non-tissue spots. Datasets with limited background area, inaccurate tissue masks, or unusually low non-tissue signal may provide weak anchors for initialization, potentially leading to less stable early training dynamics. Third, the current implementation applies a shared diffusion-kernel structure across genes; while this is supported by our diffusion-atlas observations in many samples, gene-specific deviations can still occur in some biological contexts.

These limitations motivate several directions for future work. One possible direction is adaptive prior weighting, where image-label supervision strength is learned or uncertainty-weighted instead of fixed, allowing the model to downweight unreliable histology cues. For datasets with sparse or unreliable non-tissue spots, diffusion-rate initialization could be achieved by user defined markers for each image niche. Another extension is hierarchical or partially gene-specific diffusion kernels that preserve computational efficiency while capturing residual gene-level heterogeneity. It will also be valuable to benchmark CLEAR-ST across additional platforms, permeabilization protocols, and tissue morphologies, and perhaps to integrate richer spatial covariates (e.g., nuclei density or multiplex imaging channels) into the generative model.

In summary, CLEAR-ST provides an interpretable and practically useful approach to diffusion-aware correction in sequencing-based spatial transcriptomics. By explicitly modeling transcript spread while preserving biological structure, it improves multiple downstream analyses and offers a strong foundation for future methodological development.

## 4 Methods

### 4.1 CLEAR-ST Structure

At its core, CLEAR-ST couples a denoising autoencoder with a forward graph-based diffusion process. This unified framework is trained end-to-end via stochastic variational inference to estimate an underlying clean gene expression field from the contaminated spatial transcriptomic counts.

The denoising autoencoder employs a 2-layer multilayer perceptron (MLP) for both the encoder and decoder, with a user-defined latent dimension *d* and a dropout rate of 0.1 applied to the hidden layer to prevent overfitting. Let **Y ∈** R*^N×G^* represent the raw count matrix for *N* spots and *G* genes. The log-transformed counts are first passed through the encoder network, consisting of linear transformations followed by Layer Normalization, GELU activation, and dropout:

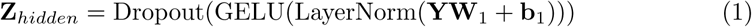

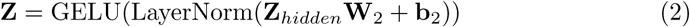

where **Z ∈** R*^N×d^* is a latent representation capturing the underlying expression structure. The decoder network then projects **Z** back to the log-scale gene expression space, **X ∈** R*^N×G^*, using a symmetric architecture. CLEAR-ST uses image-derived histological niche labels as an auxiliary regularization prior, as histology provides complementary structural information that is not directly affected by transcript capture artifacts. A classifier head built on top of the latent embedding **Z** predicts the pre-assigned niche labels, generating an auxiliary loss weighted by a user-adjustable parameter *λ*_niche_, encouraging the latent representation to retain structural consistency with image-derived niches without enforcing hard boundaries.

To model the physical process of mRNA lateral diffusion, CLEAR-ST models contamination using a discrete approximation of the heat equation over the tissue space. In classical physics, diffusion of a continuous quantity is commonly described by the heat equation. Discretizing this spatial spread over the expression spots yields a graph-Laplacian-based forward diffusion operator.

To account for structural biases – where mRNA might diffuse more freely along specific tissue axes as we have shown – the spatial adjacency graph is constructed using a non-elliptical rotated diffusion kernel. Let (*x_i_, y_i_*) denote the spatial coordinates of spot *i*, and let *θ* be a global orientation angle. For each directed edge from spot *i* to spot *j*, we define the raw coordinate differences Δ*x* = *x_j_ - x_i_* and Δ*y* = *y_j_ - y_i_*, and compute the rotated coordinates:

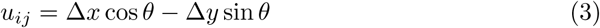

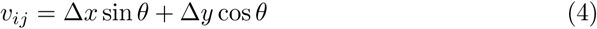

To allow the elongation of the kernel to differ between the positive and negative directions along each rotated axis, we split *u_ij_* and *v_ij_* into their positive and negative rectified components, *u^+^_ij_ = max(u_ij_,0), u^−^_ij_ = max(−u_ij_,0), v^+^_ij_ = max(v_ij_,0), v^−^_ij_ = max(−v_ij_,0)*. The directed, non-elliptical squared distance is:

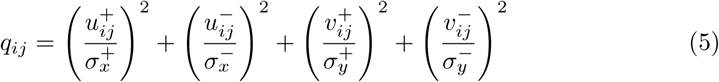

Where *σ^+^_x_, σ^−^_x_, σ^+^_y_, σ^−^_y_* are direction-specific principal diffusion lengths, each controlling the kernel spread in one of four signed directions in the rotated frame. The edge weight is then computed via a Gaussian kernel with an additional learnable sharpness parameter *τ* that modulates the decay rate:

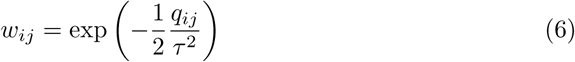

After computing all edge weights, we apply row-wise normalization so that for each source spot *i*, the outgoing weights sum to *1: w~_ij_ = w_ij_/Σ_k_w_ik_*, ensuring that the diffusion operator conserves total mass. During training, an entropy penalty *−λ_H_Σ_i_Σ_j_w~_ij_logw~_ij_* (with small *λ_H_* = 10*^−^*^3^) is applied to the outgoing weight distribution of each spot, encouraging the kernel to distribute mass across multiple neighbors rather than collapsing to a single direction. By learning the four directional sigmas, the global rotation *θ*, and the sharpness *τ*, this non-elliptical formulation allows the model to represent asymmetric, directionally biased diffusion: mRNA leakage may spread farther in one direction than its opposite, a pattern we observed empirically in real samples.

#### 4.1.1 Physical interpretation of the weight matrix and forward diffusion

Having defined the normalized edge weights, we now clarify the physical meaning and orientation of the resulting weight matrix **W**, whose entries are *w*~*_ij_* (we drop the tilde in subsequent notation for brevity). Each entry *w_ij_* quantifies the fraction of transcript mass that moves from spot *i* to spot *j* in one discrete diffusion step. **W** is therefore a row-stochastic transition matrix (*Σ_j_w_ij_ = 1* for every source spot *i*), which guarantees that the total transcript count is conserved during redistribution—transcripts are neither created nor destroyed, only relocated among spots.

The orientation convention *w_ij_* = flow from *i* to *j* is chosen to match the physical process of the diffusion process during experimentation. In a capture-based spatial transcriptomics assay, transcripts are released from their cells of origin during permeabilization, diffuse through the tissue section, and bind to oligonucleotide probes coating each spot. The clean expression field **X** represents the transcript abundance at each source location *before* diffusion, and applying **W** from the right as **XW**^T^ (or equivalently, the graph Laplacian **L** as **LX** in the update below) propagates mass *forward* from sources to their neighbors, yielding the expected contaminated observation.

We deliberately model the *forward* process rather than attempting to directly invert the contamination. The forward model is well-posed: a given clean field, together with specified diffusion parameters, uniquely determines the expected contaminated field. The inverse problem—recovering the clean field from contaminated observations—is ill-posed, because multiple distinct clean configurations can yield indistinguishable contaminated observations after spatial mixing. CLEAR-ST resolves this ambiguity by coupling the forward diffusion operator with a denoising autoencoder, which provides a learned prior over plausible clean expression patterns. Joint optimization under the probabilistic observation model then identifies the clean-field estimate that, when passed through the forward process, best explains the observed counts.

#### 4.1.2 Graph heat equation and forward update

In the classical heat equation on a continuous domain, the rate of change of a signal (e.g., temperature or concentration) at a point is determined by the difference between that point and its spatial neighborhood. According to Newton’s law of cooling, heat flows from regions of higher concentration to nearby regions of lower concentration, and the Laplace operator captures local differences between neighboring locations [15]. On a graph, the same idea can be formulated in discrete form by replacing continuous space with a set of nodes connected by weighted edges. Suppose signals on the graph is represented by a vector **f** R*^N^*, where *f_i_* denotes the signal value at node *i*, and diffusion is driven by differences between each node and its neighbors. For a weighted graph, the combinatorial graph Laplacian is defined as

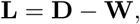

where **W** is the weighted adjacency matrix with entries *w_ij_* and **D** is the diagonal degree matrix with *D_ii_ = Σ_j_w_ij_* [15]. Applied to a graph signal, the Laplacian acts nodewise as

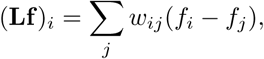

which measures how much the value at node *i* differs from the weighted average of its neighbors. As a result, the graph heat equation

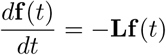

describes diffusion on the graph. In our model, we generalize this update with spot-specific diffusion rates and a small damping (teleport) term that prevents unbounded growth of the diffused field. Starting from the clean expression **X**^(0)^ = **X**, the contaminated profile after *t* steps evolves as:

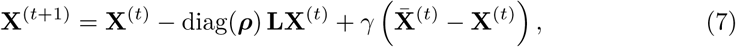

where ***ρ*** *∈* R*^N^* represents spot-specific per-step diffusion rates parameterizing local tissue permeability, *X^(t)^* is the per-gene mean expression at step *t*, and *γ ∈* [0, 1] is a small teleport parameter (default 0.01, constrained to [0, 0.05]). This update is applied for *T* steps (default *T* = 10, configurable by the user, using 5-10 steps is generally appropriate) to produce the final contaminated expression *X~ = X^(T)^*. The resulting diffused expression represents the expected contaminated observation under the forward model.

The diffused continuous prediction X~ is linked to the raw integer observations **Y** through an explicit probabilistic count model. By default, CLEAR-ST uses a Zero-inflated Poisson (ZIP) likelihood, which captures both the count nature of the data and accommodates zero-inflation commonly observed in spatial transcriptomics, while maintaining a relatively simple parameterization compared to more flexible count models such as the Zero-inflated Negative Binomial (ZINB). Other likelihood models including ZINB, Negative Binomial (NB), Poisson, Gamma Poisson (GP) are also available for users to try. Model parameters were optimized using stochastic variational inference with the Adam optimizer (default learning rate 1 10*^−^*^3^). Training was carried out for 500–1000 epochs. Convergence was assessed based on stabilization of the evidence lower bound (ELBO).

### 4.2 Parameter Initialization

Initialization of spatial image labels and expression-based clustering labels provides useful starting points for separating structured biological variation from diffusion-associated artifacts. The former acts as the regularization force from image-derived niches (an auxiliary source of structural information) and the latter informs diffusion rates based on observed diffused patterns that associate out-of-tissue expression with transcriptionally similar in-tissue regions (co-clustered in-tissue spots).

Prior to model training, the spatial connectivity and distance graphs are built using the physical coordinates of the spots. High-resolution histology image patches are extracted around each spot and clustered using Leiden to define histology-derived *niche labels*. Similarly, the expression profiles are clustered to derive expression-based labels based on standard preprocessing after the standard preprocessing pipeline, which link transcriptionally similar out-of-tissue spots to in-tissue spots. The optimal clustering is automatically selected by maximizing the silhouette score as a heuristic criterion, with a minimum cluster number of 3 in both cases.

The parameters defining the diffusion operator are initialized based on these structural relationships prior to variational inference. To initialize the spot-specific diffusion rates, CLEAR-ST first estimates a global contamination level using the proportion of signal detected in background (non-tissue) spots as a proxy. We employ a niche-based aggregation approach: for each expression-derived niche *k*, the model aggregates transcript counts within in-tissue spots assigned to each expression-derived niche, *C*_tissue*,k*_, and the background counts assigned with the same expression-derived niche, *C*_bg*,k*_. The total diffusion fraction for niche *k* is calculated as the corresponding contamination fraction for niche *k* is defined as the ratio of background-associated counts to total counts within that niche (*C_k_* = *C*_bg*,k*_*/*(*C*_tissue*,k*_ + *C*_bg*,k*_)) and clamped between 0.1% and 1.5× the global contamination percentage to avoid extreme parameter values during initialization. Assuming *T* discrete forward diffusion steps, the initialization value for the per-step diffusion-rate parameter for niche *k* is 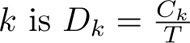. Each spot *i* is then assigned its initial rate *ρ^(0)^_i_* based on its niche label. During the probabilistic inference phase, these spot-specific diffusion rates are modeled using beta distributions, enabling *ρ_i_* to progressively diverge from the niche-level initialization and vary independently, thereby reflecting local variation in diffusion-related effects.

Furthermore, to accommodate directional leakage, we defined parameters of the non-elliptical diffusion kernel – namely the direction-specific principal diffusion lengths *σ^+^_x_, σ^−^_x_, σ^+^_y_, σ^−^_y_*, the rotation angle (*θ*), and the sharpness parameter (*τ*). To provide a physically meaningful scale, all four directional spread widths are initialized to the median Euclidean distance between adjacent spots in the nearest-neighbor connectivity graph (*σ*_init_ = median(*d_ij_*)). This establishes a sensible null prior that baseline leakage primarily impacts the immediate 1–2 spot grid neighborhood. *θ* is initialized at 0.1 radians (near zero) to start with a weakly rotated kernel, and *τ* is initialized at 1.0, corresponding to a standard Gaussian decay without additional sharpening or flattening. Combined, these parameter initializations ensure CLEAR-ST initiates diffusion on an anatomically constrained scale before allowing the data to refine the precise directional widths, angular orientation, and decay sharpness during SVI training.

### 4.3 Diffusion Investigation Metrics and Tests

To thoroughly characterize the mRNA spatial diffusion artifact, we developed a suite of metrics quantifying the extent, spatial decay, and directional coherence (anisotropy) of the out-of-tissue expression.

#### 4.3.1 Out-of-Tissue Fraction and Spatial Decay

We quantified the global diffusion burden for each dataset using the out-of-tissue expression fraction, defined as the ratio of total counts in background (out-of-tissue) spots to the total counts across the entire slide. To model how leaked signal decays away from the tissue boundary, we mapped each out-of-tissue spot to its nearest in-tissue spot using a KD-Tree to calculate the Euclidean distance *d_i_* to the tissue edge. For individual genes, we constructed distance-decay profiles by calculating the mean normalized expression in concentric bins and fitted a log-linear exponential decay model:

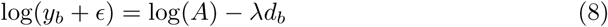

where *y_b_* is the mean expression in distance bin *b*, *d_b_* is the distance to the bin center, *λ* is the decay constant, and *A* estimates the theoretical expression at the boundary edge. Larger *λ* values indicate steeper decay.

#### 4.3.2 Directional Anisotropy of Diffusion

Physical permeabilization and fluid flow can induce directional leakage. We quantified diffusion anisotropy per gene per sample by computing the resultant vector length **R** from out-of-tissue spots. For each gene *g*, let *w_i_* be its expression in an out-of-tissue spot *i*, and *θ_i_* be the polar angle of the vector pointing from its nearest in-tissue spot to spot *i*. Specifically, the direction vector for each out-spot *i* was Δ**v***_i_* = (*x_i_ − x_n_*_(*i*)_*, y_i_ − y_n_*_(*i*)_), where *n*(*i*) is the nearest in-tissue spot. The polar angle was computed as *θ_i_* = atan2(Δ*y_i_,* Δ*x_i_*), with 0 radians along the +*x* (right/east) axis and positive angles counter-clockwise. The resultant vector components were computed as *C = Σ_i_w_i_cos θ_i_* and *S = Σ_i_w_i_sin θ_i_*. The anisotropy magnitude was defined as:

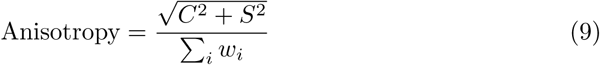

Values closer to 1 indicate highly unidirectional leakage, while values near 0 indicate isotropic circular diffusion. The preferred diffusion direction was computed as the phase angle *θ*_pref_ = atan2(*S, C*), which can be interpreted as the dominant leakage direction of this gene.

We summarized directional heterogeneity across genes within each slide by computing the Shannon entropy of the distribution of preferred angles. Let *θ*_pref*,g*_ be the preferred direction for gene *g* in a slide. We binned *θ*_pref*,g*_ into *B* equal-width bins over [*−π, π*),yielding counts *h_b_* and probabilities *p_b_ = h_b_/Σ^B^_b=1_h_b_*. Orientation entropy was then:

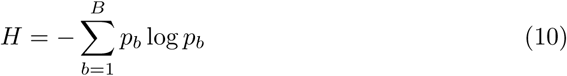

Higher *H* indicates more uniformly spread directions and lower *H* indicates directional concentration.

For visualization, we plotted a polar “compass” by placing each angular bin’s center at its corresponding angle and using the bin height to encode its frequency. Specifically, the compass curve uses (*θ_b_, p_b_*), where *θ_b_* is the center angle of bin *b* and the radial coordinate is the normalized frequency *p_b_*. This renders the preferred-direction distribution as a radial profile around the circle.

#### 4.3.3 Binomial Regression for Gene Determinants

To identify intrinsic and spatial determinants of a gene’s diffusion propensity, we modeled the proportion of out-of-tissue counts over total counts using a binomial generalized linear mixed model (GLM) with a logit link. The dataset ID was included as a fixed effect. The base probability of out-of-tissue placement for gene *g* in dataset *j*, *p_g,j_*, was parameterized as:

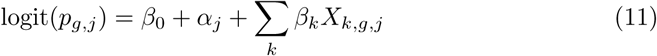

where *α_j_*is the dataset-specific intercept, and *X_k,g,j_* are covariates including log-transformed total gene counts, the variance-to-mean ratio computed on in-tissue spots *(log(CV^2^_in_))*, fractional tissue detection rate, number of in-tissue spots, mitochondrial gene indicator, and log-transformed exon union length.

### 4.4 Benchmarking and Downstream Analyses

To assess the effectiveness of CLEAR-ST in correcting spatial transcriptomics data, we developed an evaluation pipeline benchmarking the corrected output against the original uncorrected data and alternative established methods (e.g., SpotClean, resolVI). Prior to downstream benchmarking, uncorrected and corrected count matrices undergo standard single-cell preprocessing utilizing the scanpy library [16]. Total counts are normalized to 10^4^ per spot, followed by log(1 + *x*) transformation. Highly variable genes are identified using the Seurat v3 dispersion-based method. Finally, data is scaled to zero mean and unit variance before running Principal Component Analysis (PCA) to calculate lower-dimensional embeddings and nearest-neighbor graphs.

#### 4.4.1 Clustering and Spatial Autocorrelation Evaluation

To evaluate the impact of diffusion correction on spatial structure, we compute Moran’s I spatial autocorrelation for all genes using squidpy [17]. An increase in Moran’s I indicates enhanced spatial coherence of gene expression following correction. For spatial clustering evaluation, we apply Leiden clustering to the precomputed nearest-neighbor graphs. We quantify clustering consistency and agreement using standard metrics from the scikit-learn library [18], including Silhouette Score for intra-cluster cohesion and inter-cluster separation based on PCA embeddings, NMI for agreement between clustering assignments adjusting for chance, and Homogeneity and Completeness for clusters contain only spots of a single class and that spots of a given class are assigned to the same cluster.

#### 4.4.2 Markers and GSEA

We first identify corresponding pre- and post-correction clusters using a custom overlap matching procedure. Let *S^bf^_u_* and *S^af^_v_* denote spot sets of cluster *u* (before) and cluster *v* (after). Their overlap is computed by the Jaccard index

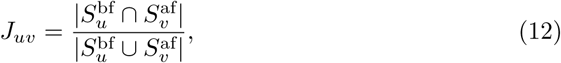

and a greedy one-to-one matching is performed by repeatedly selecting the largest remaining *J_uv_*, with a minimum threshold of 0.6. This yields matched cluster pairs and unmatched clusters for stability assessment.

Marker genes are then computed separately for before and after objects with `scanpy.tl.rank genes groups (Wilcoxon test, use raw=True, pts=True)`. We retain positive and confident markers using: adjusted *p*-value *<* 0.05, logFC *>* 0, and in-cluster detection rate `pct nz group` *>* 0.1. For each matched cluster pair, we compare (i) the number of significant markers, (ii) the overlap of marker sets, and (iii) effect-size shifts of shared markers using

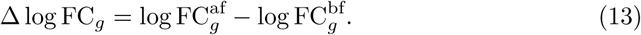

For pathway-level interpretation, we run GSEA with gseapy under two designs: matched-cluster before-vs-after contrasts and annotation-stratified before-vs-after contrasts. We call gp.gsea (the expression based method) with phenotype permutations (*n* = 1000), and gene sets from GO Biological Process 2023, MSigDB Hallmark 2020, and Reactome 2022 (default: min size=5, max size=2000). Pathways are summarized by normalized enrichment score (NES) and FDR *q*-value; significance is defined as FDR *<* 0.05, and direction is assigned by the sign of NES (up-regulated if NES*>* 0, down-regulated if NES*<* 0).

#### 4.4.3 Cell2location model fitting settings

For each slide, Cell2location was run with the same pipeline and hyperparameters across for original and CLEAR-ST counts. Briefly, we first fit the reference Regression-Model using cell type level 3 labels and donor age as batch covariate, with gene filtering thresholds cell count cutoff = 5, cell percentage cutoff2 = 0.03, and nonz mean cutoff = 1.12, for 2000 epochs (batch size 3000). We then intersected genes between reference signatures and spatial data, removed mitochondrial genes, and fit the spatial Cell2location model with N cells per location = 10 and detection alpha = 20 for 3000 training epochs. Posterior summaries were exported using 1000 posterior samples (batch size equal to the number of spots), and the inferred abundance matrices and posterior quantiles were used for downstream analyses.

#### 4.4.4 Run Benchmarking Methods

We compared CLEAR-ST against two alternative methods designed to address similar issues. For both SpotClean and resolVI, we ran them on the same highly variable expression subset, following their recommended workflow. For resolVI, we used the unsupervised mode and ran for 400 epochs. We then sampled from posterior to generate the corrected counts for downstream analyses and evaluation. For SpotClean, we used the recommended candidate radius of 20 and maximum iteration of 10.

## Supplementary information

Supplementary figures are provided at the end of this manuscript.

## Acknowledgements

This work was supported by the School of Biomedical Sciences, Li Ka Shing Faculty of Medicine, The University of Hong Kong, and the Laboratory of Data Discovery for Health Limited (D24H), Hong Kong Science Park.

## Declarations

- **Funding** This work was supported in part by The Hong Kong Research Grant Council General Research Fund (17123223 for JWKH), Collaborative Research Fund (C5017-24G to JWKH), Guangdong Natural Science Fund(2023A1515011265 to JWKH), NSFC/RGC Joint Research Scheme (N HKU731/21 to JWKH), the Shenzhen-Hong Kong-Macau Technology Research Programme (Type C; SGDX2021082310356025 to JWKH), and the AIR@InnoHK programme by the Innovation and Technology Commission of Hong Kong.
- **Conflict of interest/Competing interests** No competing interest is declared.
- **Ethics approval and consent to participate** Not applicable.
- **Consent for publication** Not applicable.
- **Data availability** All datasets used and mentioned are available publically. The Valdeolivas 2024 dataset is deposited at https://zenodo.org/records/7760264, with the processed and labeled objects stored at https://cellxgene.cziscience.com/collections/68cba939-4e72-4405-80ef-512a05044fba. The 10X datasets are downloaded from https://www.10xgenomics.com/datasets. The Yayon 2024 dataset has raw data deposited at https://www.ebi.ac.uk/ena/browser/view/PRJEB77091, and processed data with labels at https://cellxgene.cziscience.com/collections/fc19ae6c-d7c1-4dce-b703-62c5d52061b4.
- **Materials availability** Not applicable.
- **Code availability** CLEAR-ST is available on GitHub at https://github.com/holab-hku/CLEAR-ST.
- **Author contribution** K.M. developed the method, performed the analyses, and wrote the manuscript. Y.H. and J.W.K.H. supervised the study and revised the manuscript. These authors contributed equally: Kun Ma.

