## Supplementary Figure 1-3 for "CLEAR-ST: Physics-informed probabilistic decontamination of spatial transcriptomics by modeling mRNA lateral diffusion"

### Supplementary Figures

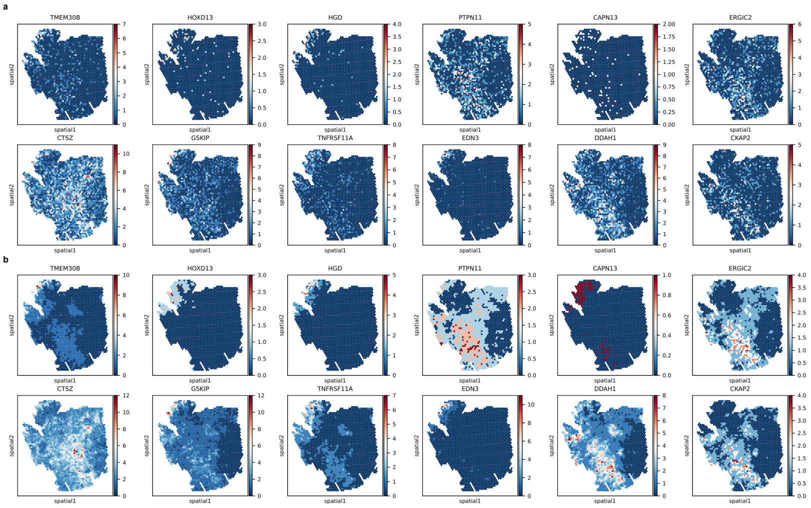


Supplementary Fig. 1: For sample `S6_Rec_A938797_Rep1`, spatial patterns of genes with top Moran's I improvements before (**a**) and after correction (**b**).

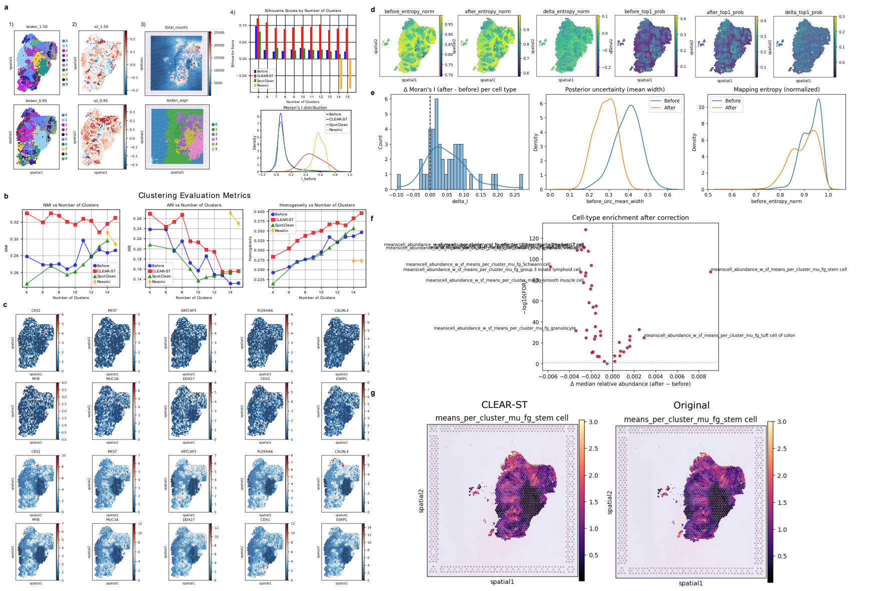


Supplementary Fig. 2: CLEAR-ST corrected expression evaluation for sample `S6_Rec_A938797_Rep1`. **a,** (1) Spatial cluster label assignment at a higher matching cluster number for before (top row) and after (bottom row) CLEAR-ST correction and (2) the spot-level silhouette scores. (3) Total counts on the whole slide and the co-clustering pattern of in-tissue and out-of-tissue spots. (4) Total silhouette scores across cluster numbers and Moran's I distribution. **b,** Clustering evaluation metrics across cluster numbers for each evaluation method. **c,** Spatial expression of genes with the largest improvements in Moran's I before (top two rows) and after correction (bottom two rows). **d,** Single-cell reference mapping entropy for before and after correction, changes in cell type entropy (mostly negative), top 1 cell type assignment before and after correction, and changes in top 1 cell type relative abundance. **e,** Changes in Moran's I for cell type abundances, posterior mean width distribution for before and after correction, and entropy distribution for before and after. **f,** Volcano plot showing the most enrichment-enhanced cell types and the most decreased cell types. **g,** Spatial distribution of stem cell abundance before (left) and after correction (right). The improvement mainly took place in the tumor and `stroma_fibroblastic_IC_high` regions.

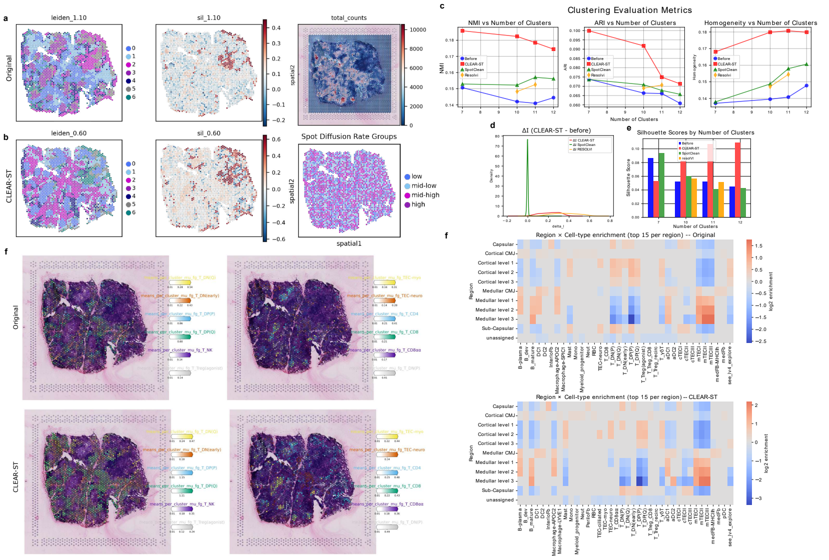


Supplementary Fig. 3: Corrected counts evaluation results of sample TA11486163. For a matching cluster number $k = 7$, the raw (**a**) and corrected (**b**) count-derived niche labels and spot-level silhouette scores. **c,** Clustering evaluation metrics across benchmarking methods. **d,** Changes in gene Moran's I. **e,** Silhouette scores at the same cluster numbers across correction methods. **f,** Full region × cell-type enrichment heatmap for before (top) and after correction (bottom). **g,** Simultaneous cell-type visualization showing the enrichment of T DP/DN subtypes.
